# Identification of new *Dickeya dadantii* virulence factors secreted by the type 2 secretion system

**DOI:** 10.1101/2021.08.03.454866

**Authors:** Guy Condemine, Bastien Le Derout

**Affiliations:** Univ Lyon, Université Lyon 1, INSA de Lyon, CNRS UMR 5240 Microbiologie Adaptation et Pathogénie, F-69622 Villeurbanne, France

## Abstract

*Dickeya* are plant pathogenic bacteria able to provoke disease on a wide range of plants. A type 2 secretion system (T2SS) named Out is necessary for bacterial virulence. Its study in *D. dadantii* showed that it secretes a wide range of plant cell wall degrading enzymes, including pectinases and a cellulase. However, the full repertoire of exoproteins it can secrete has probably not yet been identified. Secreted proteins are first addressed to the periplasm before their secretion by Out. No secretion signal present on the protein allows the identification of substrates of a T2SS. To identify new Out substrates, we analyzed *D. dadantii* transcriptome data obtained in plant infection condition and searched for genes strongly induced encoding a protein with a signal sequence. We identified four new Out-secreted proteins: the expansin YoaJ, the putative virulence factor VirK and two proteins of the DUF 4879 family, SvfA and SvfB. We showed that SvfA and SvfB are required for full virulence of *D. dadantii* and showed that *svf* genes are present in a variable number of copies in other *Pectobacteriaceae*, up to three in *D. fanghzongdai*. This work opens the way to the study of the role of non-pectinolytic proteins secreted by the Out pathway in *Pectobacteriaceae*.

**IMPORTANCE:** The plant pathogen *Dickeya* rely on a type 2 secretion system named Out for their pathogenicity. Importance of plant cell wall degrading enzymes secreted by this system has been well studied. However, existence and role of other Out-secreted proteins has barely been investigated. By mining *D. dadantii* transcriptome data, we identified four new Out-secreted proteins. We showed that two of them, SvfA and SvfB, are necessary for the full virulence of the bacteria. These findings show that identification of all the proteins secreted by the *Dickeya* Out system is necessary for a better knowledge of the virulence of these bacteria.

## INTRODUCTION

Soft rot *Pectobacteriaceae* (SRP), *Dickeya* and *Pectobacterium*, are plant pathogenic bacteria that can provoke disease on more than 35% of angiosperm plant orders, including both monocot and dicot plants (1). Among those, there is a wide range of plants of agronomic interest such as potato, rice, chicory, cabbage or ornementals on which they can cause severe losses. Symptoms are usually soft rot but these bacteria can provoke blackleg or wilting on aerial parts of potato. Recently, diseases on woody plants caused by *Dickeya* have been reported (2). There is no efficient way to fight these bacterial diseases. There are actually twelve species of *Dickeya* described, isolated either from infected plants (type strain of *D. chrysanthemi* isolated from *Chrysanthemum morifolium*, *D. dadantii* subsp. *dadantii* from *Pelargonium capitum*, *D. dadantii* subsp. *diffenbachiae* from *Dieffenbachia* sp., *D. dianthicola* from *Dianthus caryophillus*, *D. zeae* from *Zea mays*, *D. oryzae* from *Oryza sativa*, *D. paradisiaca* from *Musa paradisiaca*, *D. solani* from *Solanum tuberosum*, *D. fangzhongdai* from *Pyrus pyrifolia*, *D. poaceiphila* from *Saccharum officinarum*) (3)(4)(5)(6)(7) or from river of lake waters (*D. aquatica*, *D. lacustris* and *D. undicola*) (8)(9)(10). The role of protein secretion systems on the onset of the disease provoked by these bacteria has been recognized long ago (11). In contrast to many plant pathogenic bacteria, the type three Hrp secretion system is not the main determinant for SRP virulence (12). The main virulence factor for these bacteria is a type 2 secretion system (T2SS) named Out. It allows the secretion of enzymes that degrade the components of the plant cell wall, leading to the soft rot symptom distinctive of the disease. The first Out-secreted proteins to be identified were a set of pectinases and a cellulase which are easily detectable by simple enzymatic tests (13) (11). The pectinolytic secretome of the model strain *D. dadantii* 3937 has been studied in detail by cloning the genes of these easily detectable enzymes. *D. dadantii* secretes by the Out machinery nine pectate lyases, one pectin methylesterase, one pectin acetylesterase and one rhamnogalacturonate lyase (14). A proteomic analysis of the secreted proteins by 2D gel electrophoresis allowed the identification of two other secreted proteins, the feruloyl esterase FaeD and a protein with homology with a *Xanthomonas campestris* avirulence protein AvrL (15). A search in *D. dadantii* of homologues of proteins secreted by the Out T2SS of *Pectobacterium atrosepticum* (16) recently led to the characterization of the metal binding protein IbpS (17). There is no strict host specificity for Dickeya species, however some of them show a preference for some plant species. Since all the pectinolytic enzymes studied in *D. dadantii* are present in most of other *Dickeya* species these enzymes are probably not responsible for the host preference observed for these bacteria (18). We hypothesized that additional T2SS-secreted proteins specific for some species might exist and play a role in the host preference. To identify such proteins, we analyzed previously published *D. dadantii* transcriptome data, looking for genes induced in plant infection conditions and encoding proteins with a signal sequence. We identified several proteins secreted by the Out machinery and showed that two proteins of the DUF4879 family, SvfA and SvfB are *D. dadantii* virulence factors.

## MATERIAL AND METHODS

### Bacterial strains and growth conditions

Bacterial strains, phages, plasmids and oligonucleotides used in this study are described in Table 2. *D. dadantii* and *E. coli* cells were grown at 30 and 37°C respectively in LB medium or M63 minimal medium supplemented with a carbon source (0.2%, w/v unless otherwise indicated). When required antibiotics were added at the following concentrations: ampicillin, 100 mg l^-1^, kanamycin and chloramphenicol, 25 mg l^-1^. Media were solidified with 1.5% (w/v) agar. Transduction with phage ΦEC2 was performed according to Résibois *et al*. (40)

### Mutant construction

To construct strain A6418 that contains a *svfA-uidA* fusion a 1.3 kb DNA fragment containing *svfA* was amplified with primers 17176H+ and 17176A. The resulting fragment was inserted into the pGEM-T plasmid (Promega). A XbaI site was created by site directed mutagenesis with the primers 17176XbaF and 17176XbaR into the *svfA* coding sequence and a *uidA*-kanR cassette was inserted into this XbaI site. To construct strain A6467 that contains a *svfB-uidA-kanR* fusion and a CmR cassette a 2000bp DNA fragment containing *svfB* was amplified with the primers 15544L2+ and 15544L2-. The resulting fragment was inserted into the pGEM-T plasmid. A XmaI site was created by site directed mutagenesis into *svfB* coding sequence with the primers 15544XmaF and 15544XmaR and a *uidA*-kanR cassette was inserted into this created unique XmaI site. All the constructs were recombined into the *D. dadantii* chromosome according to Roeder and Collmer (41). Recombinations were checked by PCR. His-tagged versions of the proteins SvfA, SvfB, YoaJ and VirK were constructed by amplifying the corresponding genes with the primers 17176H+ and 17176H-, 15544H+ and 15544H-, 14642H+ and 14642H-, VirKH+ and VirKH-, respectively. The resulting DNA fragments were cloned into plasmid pGEMT.

### Secretion assays and Western blots

*D. dadantii* strains containing the plasmid to test were grown overnight in LB medium in the presence of the appropriate antibiotic. Culture supernatant containing secreted proteins was separated from cells by centrifugation at 10,000 g for 3 min, and both fractions were loaded onto 12% polyacrylamide gel electrophoresis (SDS-PAGE). The proteins were next transferred onto Immobilon P membrane (Merck) and probed with Ni-NTA-HRP.

### Pathogenicity tests

Bacteria were grown overnight in LB medium, centrifuged and resuspended at OD_600_ 1 in M63 medium. Potatoes were surface sterilized with 70% ethanol and dried. A hole was made with a pipette tip and 10 *μ*l of bacteria were deposited in the hole which was covered with mineral oil. Potatoes were placed over a wet paper in a tray contained in a plastic bag to maintain moisture. After 48 h at 30°C, the weight of rotten tissue was measured.

### Enzymatic assays

β-glucuronidase assays were performed on toluenized extracts of cells grown to exponential phase using the method of Bardonnet *et al* (42) with *p*-nitrophenyl-β-D-glucuronate as the substrate.

## RESULTS

### Identification of new Out-secreted proteins

To have a more complete knowledge of the proteins secreted by the *D. dadantii* Out T2SS that could be involved in the pathogenicity process, we searched for candidate genes in recently published transcriptome data (19)(20). We selected genes strongly induced during plant infection and coding for a protein possessing a signal sequence. We retained the genes *Dda3937_01687*, *Dda3937_00585* (thereafter named SvfA and SvfB, respectively) and *Dda3937_00081* (also named *yoaJ*). We also retained VirK, a protein of unknown function with a signal sequence identified among the genes controlled by the transcriptional regulator PecS of many virulence factors (21). Each protein was tagged with a His-tag and its secretion was analyzed in the *D. dadantii* wild type strain and an *outD* mutant in which the Out machinery is not functional. The proteins SvfA, SvfB, YoaJ and VirK were detected in the supernatant of the wild type strain but not of the mutant, demonstrating their secretion by the Out machinery (Fig. 1). Production of these proteins from a gene cloned on a multicopy plasmid may explain why secretion was not total in the wild type strain.

**Fig 1:**
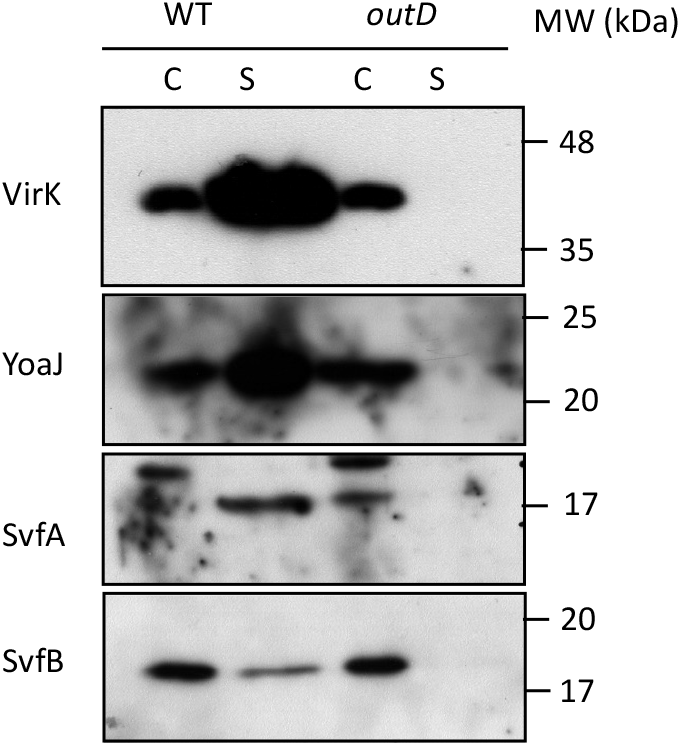
Identification of new secreted proteins by *D. dadantii*. Wild-type and *outD* mutant strains containing plasmid bearing the gene of the selected protein were grown overnight in LB medium. The supernatant (S) and cellular (C) fractions were separated by SDS-PAGE. After blotting, the proteins were detected with Ni-NTA-HRP.

YoaJ is a PecS-regulated gene (21) and it was found among the most induced genes during *Arabidopsis* infection or culture in the presence of plant extracts (19). It encodes a protein with homology to expansins. These proteins are able to non-enzymatically loosen cell wall cellulose. They are found in all plants where they have a role in cell wall extension and also in many plant pathogenic microorganisms (22). Their role in virulence has been shown in *Ralstonia solanacearum*, in *P. atrosepticum*, *P. Brasiliense* and in the plant pathogen *Erwinia tracheiphila* (23) (24). The role of *D. dadantii* expansin is probably the same. VirK is a protein of unknown function that has homologues in several plant pathogenic bacteria such as *R. solanacearum*, *Agrobacterium tumefaciens*, *Lonsdalea* and *Xanthomonas*. VirK is controlled by PecS and induced during Arabidopsis infection or culture in the presence of plant extracts (21) (20). No symptom for the *D. dadantii virK* mutant was observed whatever the plant tested (21). The two proteins SvfA and SvfB are studied in the next paragraphs.

### SvfA and SvfB are virulence factors

*svfA* and *svfB* are among the most induced *D. dadantii* genes during Arabidopsis infection or during culture of the bacteria in the presence of plant extracts (19). They are also strongly expressed during maceration of potato tubers by *D. dianthicola* and *D. solani* (25). The two *D. dadantii* proteins share 43% identity and 58% similarity in amino acid composition (Fig. 2). SvfA is 187 amino acid long (165 for the mature form, 17.5 kDa) and SvfB is 198 amino acid long (177 for the mature form, 18.9 kDa). These proteins belong to the DUF 4879 family of proteins. Proteins of this family have no known function. YolA,a protein of the DUF 4879 family showing low homology with SvfB, is among the most highly secreted protein of *Bacillus subtilis* (26). YolA is also present in *B. cereus* and in the insect pathogen *B. thuringiensis. B. subtilis* YolA is shorter than SvfA and SvfB, missing the N-terminal more variable part (Fig. 2).

**Fig. 2:**
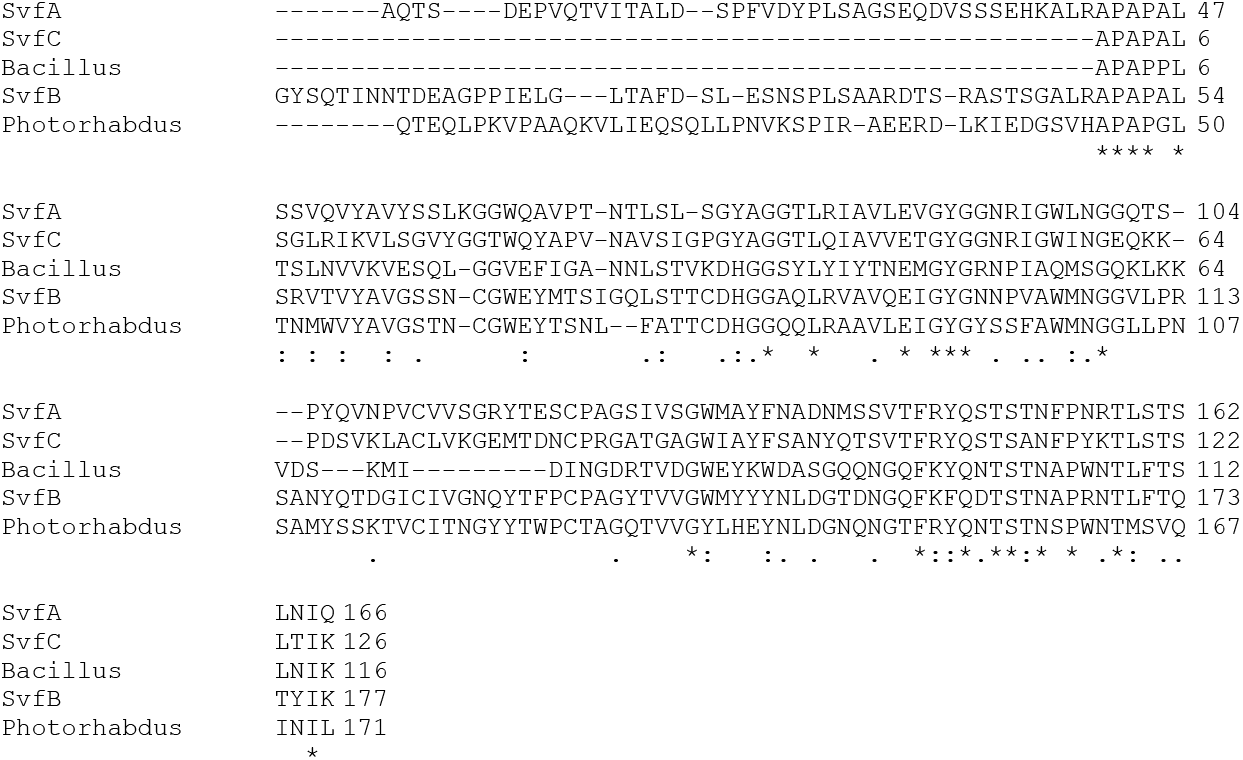
Alignment of Svf proteins. The sequences of *D. dadantii* SvfA (Dda3937_01687) and SvfB (Dda3937_00585), *D. fanghzongdai* SvfC (CVE23_15565), *B. cereus* WP-193674364.1 and *Photorhabdus asymbiotica* CAQ86327.1, without their signal sequence, were aligned with Clustal omega. Identical residues are indicated by a star and chemically equivalent residues by a double dot.

*svfA* and *svfB* mutants have been constructed and their pathogenicity has been tested on potato. The *svfA* mutant was significantly less aggressive than the wild type strain while the *svfB* mutant was not significantly affected (Fig. 3A). Virulence of the *svfA* mutant could be restored by introduction of a plasmid bearing the wild type *svfA* gene (Fig. 3B). Virulence of the double *svfA svfB* mutant was further reduced showing that the role of SvfB is additive to that of SvfA (Fig. 3A). Thus, genes *Dda3937_01687and Dda3937_00585* were named *svfA* and *svfB* for secreted virulence factor A and B.

**Fig. 3.**
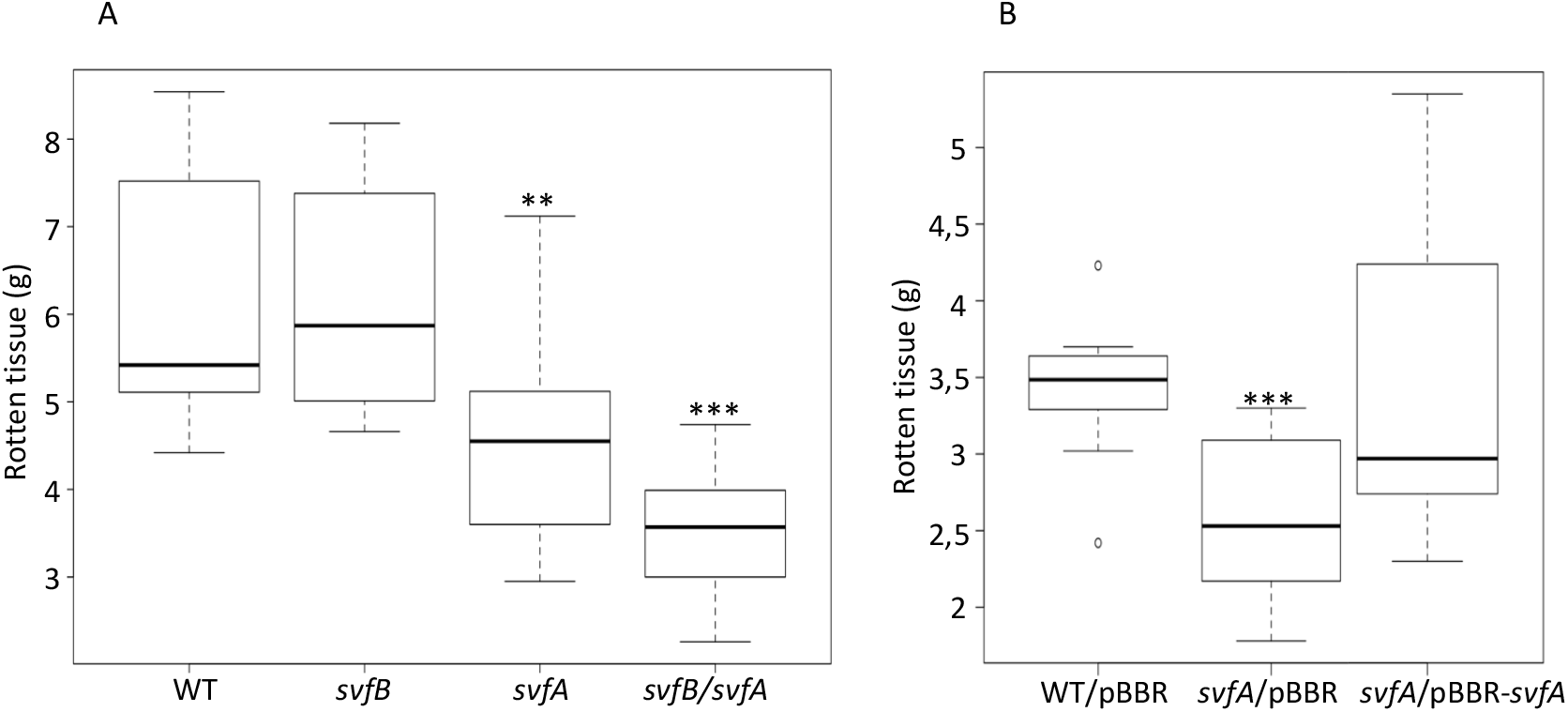
Virulence of *svfA* and *svfB* mutants. A. Potatoes were infected with the wild type strain, and the *svfA*, the *svfB* and the *svfA svfB* mutants. Rotten tissue was weighed after 48 h. B. Complementation of the *svfA* mutation. Potatoes were infected with the wild type strain, the *svfA* mutant containing the empty plasmid pBBRGm and the *svfA* mutant containing the pBBRGm plasmid bearing *svfA*. Rotten tissue was weighed after 48h. Statistical tests were performed using the Wilcoxon-Mann-Whitney test. The *p*-value were compared with an alpha risk of 4%. *p* < 0.001=***, *p* < 0.005=**, *p* < 0.01=*.

All our attempts to overproduce the proteins SvfA and SvfB in order to purify them and to study more precisely their function were unsuccessful because their production was toxic to the bacterial cells engineered to overproduce them.

### Expression of *svfA* and *svfB*

To try to identify the function of SvfA and SvfB, we analyzed the conditions in which their genes are expressed. We tested the effect of galacturonate and polygalacturonate, two compounds that are inducers of the expression of the main virulence factors, the pectate lyases, and of glucose, which represses it. We also analyzed the effect of mutations in genes controlling several aspects of *D. dadantii* virulence. KdgR represses the pectinase, pectin degradation and *out* genes (27). Its inducer is 2-keto-3-deoxygluconate, a polygalacturonate and galacturonate catabolic derivative. PecS controls genes encoding the pectinases, diverse secreted protein, the Out machinery and proteins involved in resistance to oxidative stress (28). PecT is a regulator of the pectate lyase, motility and exopolysaccharide synthesis genes (29). Pir regulates hyperinduction of pectate lyses in response to plant extracts (30). GacA, the regulator of the two-component regulatory system GacA-GacS, is a global regulator required for disease expression in response to the metabolic status of the bacteria (31). Expression of *svfA* was slightly induced by polygalacturonate but not by galacturonate (Fig. 4A). However, expression of this gene was not modified in a *kdgR* background indicating that induction by polygalacturonate is not mediated by KdgR. Growth in the presence of chicory chunks strongly induced s*vfA* expression as expected from transcriptomic data showing induction in the presence of plant extract. A high concentration of glucose led to a strong induction of *svfA* expression (Fig. 4A). This regulation is mediated by the catabolite repressor protein CRP since a mutation in the *crp* gene derepressed *svfA* expression. Thus, Crp is a repressor of *svfA*. Although it had not been previously identified as a PecS-regulated gene (21), *svf*A expression is increased in a *pecS* background. A *pir* mutation provoked a weak derepression of *svfA* expression. Neither PecT nor GacA significantly regulate *svfA* expression (Fig. 4A).

**Fig 4:**
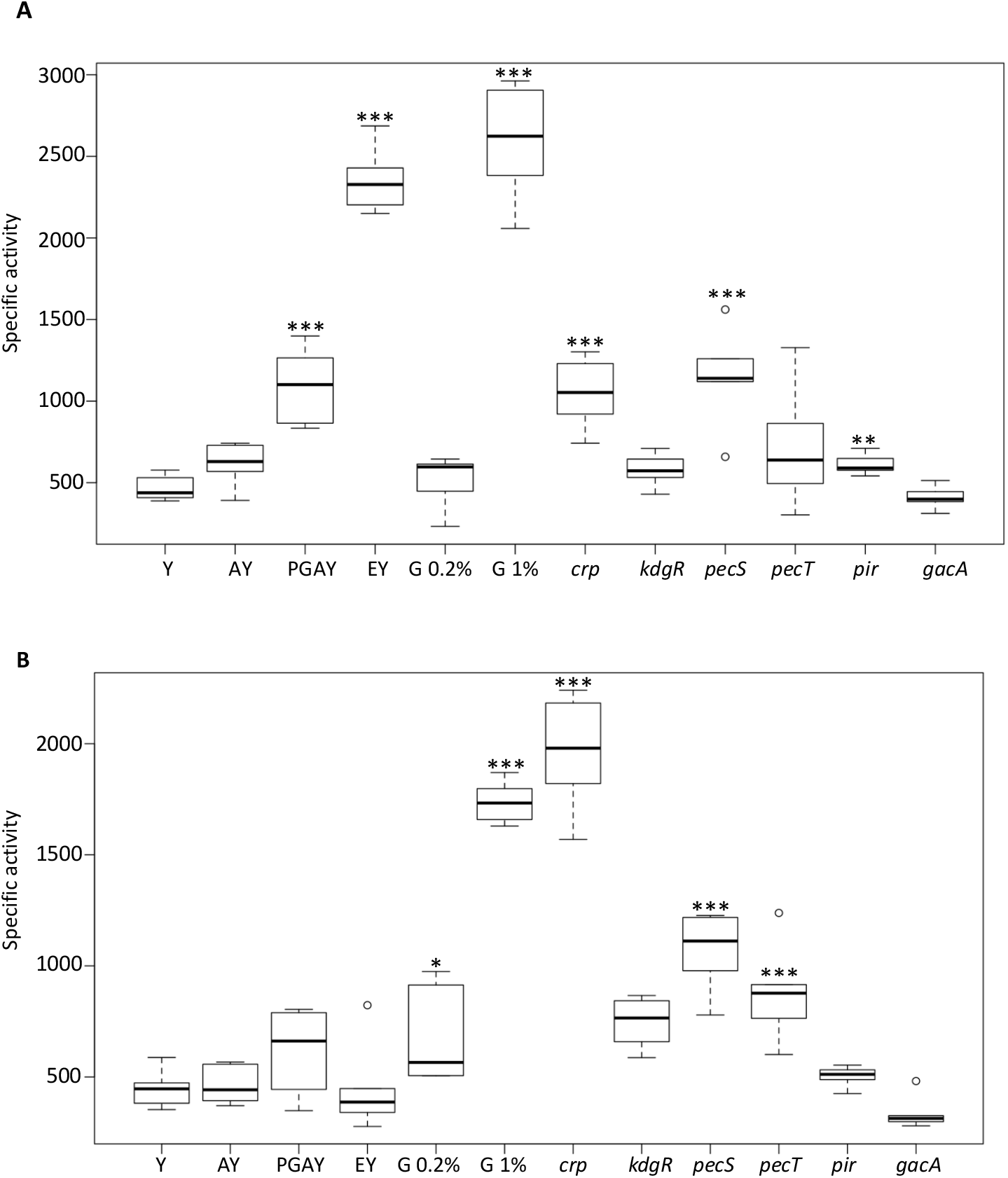
Expression of *svfA* and *svfB* in various growth conditions. A. The *D. dadantii* strain A6418 containing the *svfA-uidA* fusion and its derivative strains containing an additional regulatory mutation were grown in M63 medium in the presence of the indicated compounds (Y = glycerol, G = glucose, A = galacturonate, PGA = polygalacturonate, E = chicory chunks). Strains with additional mutations were grown with glycerol as a carbon source except the *crp* mutant that was grown with 0.2% glucose. β-glucuronidase activity was measured with *p*-nitrophenyl-β-D-glucuronate. B. Similar experiment for the *D. dadantii* strain A6467 containing the *svfB-uidA* fusion and its derivative strains containing an additional regulatory mutation. Activities are expressed in *μ*moles of *p*-nitrophenol produced per minute and per milligram of bacterial dry weight + standard deviation. Data are expressed as the mean (n = 6) from six independent experiments. Statistical tests were performed using the Wilcoxon-Mann-Whitney test. The *p*-value were compared with an alpha risk of 4%. *p* <0.001=***, *p* < 0.005=**, *p* < 0.01=*.

Regulation of *svfB* shows some similarity to that of *svfA*: it was not induced by galacturonate, polygalacturonate or regulated by KdgR, it was induced by glucose and repressed by Crp, and it was repressed by PecS (Fig. 4B). However, a few differences can be noted: in contrast to what is observed with *svfA*, no induction by plant pieces was observed for *svfB* and PecT was a repressor of *svfB* expression while Pir did not seem to control it (Fig. 4B).

### Occurrence of the new secreted proteins in other Dickeya species

Presence of *svfA*, *svfB*, *virK* and *yoaJ* was searched in the genome of all the *Dickeya* type strains, and in a few *Pectobacterium* strains (Table 1). Presence and number of proteins of the DUF 4879 family is variable among *Dickeya* species. The gene *svfA* is present in all strains except *D. zeae*, *D. chrysanthemi*, *D. poaceiphila* and *D. paradisiaca*. The gene *svfB* is present in most species but is absent in *D. chrysanthemi*, *D. poaceiphila* and *D. paradisiaca*, *D. undicola* and *D. aquatica*. A third gene located next to *svfA* and probably resulting from a duplication is found in *D. fangzhongdai* and *D. undicola*. It was named *svfC*. It has 53% homology with *D. dadantii* SvfA and 34% with *D. dadantii* SvfB. SvfC is shorter that SvfA and SvfB (126 amino acid for the mature protein, 13.2 kDa) and has the same size as *B. cereus* YolA (Fig. 2). It possesses a signal sequence, indicating that it could also be secreted by the Out system. Thus, the number of genes of the DUF 4879 family in *Dickeya* strains varies from 0 to 3. Homologues of the *svf* genes can also be found in some *Pectobacterium* strains (Table 1). For example, two copies are present in *P. carotovorum* subsp *carotovorum*. However, even in a given species, the gene may be present or not (presence of a homologue of *svfB* in 10 out of the 23 *P. brasiliense* strains present in the ASAP data bank (https://asap.ahabs.wisc.edu/asap/home.php). Outside *Pectobacteriaceae*, homologues of *svfB* can be found in a few Gammaproteobacteriaceae, i.d. in some *Photorhabdus*, *Luteibacter* and *Pseudoalteromonas* strains. *yoaJ* is present in all *Dickeya* and *Pectobacterium* strains except *D. poaceiphila* and *D. paradisiaca*. *virK* is present in all Dickeya strains except in *D. aquatica* and absent in all *Pectobacterium* strains tested.

**Table 1:** Presence of Out-secreted proteins in various *Dickeya* and *Pectobacterium* strains.

| Strain | *SvfA* | *SvfB* | *SvfC* | *YoaJ* | *VirK* | *IbpS* | *NipE* | *XynA* | *AvrL* |
| --- | --- | --- | --- | --- | --- | --- | --- | --- | --- |
| *D. dadantii* | 1 | 1 | 0 | 1 | 1 | 1 | 1 | 1 | 2 |
| *D. diffenbachiae* | 1 | 1 | 0 | 1 | 1 | 1 | 1 | 0 | 2 |
| *D. fangzhongdai* | 1 | 1 | 1 | 1 | 1 | 1 | 1 | 1 | 2 |
| *D. solani* | 1 | 1 | 0 | 1 | 1 | 1 | 1 | 1 | 2 |
| *D. zeae* | 0 | 1 | 0 | 1 | 1 | 1 | 1 | 1 | 1 |
| *D. oryzae* | 1 | 1 | 0 | 1 | 1 | 1 | 1 | 1 | 0 |
| *D. dianthicola* | 0 | 1 | 0 | 1 | 1 | 1 | 1 | 0 | 1 |
| *D. undicola* | 1 | 0 | 1 | 1 | 1 | 1 | 1 | 0 | 1 |
| *D. lacustris* | 1 | 0 | 0 | 0 | 0 | 1 | 1 | 0 | 2 |
| *D. aquatica* | 1 | 0 | 0 | 1 | 0 | 1 | 1 | 0 | 2 |
| *D. chrysanthemi* | 0 | 0 | 0 | 1 | 1 | 1 | 1 | 0 | 1 |
| *D. paradisiaca* | 0 | 0 | 0 | 0 | 1 | 0 | 0 | 0 | 0 |
| *D. poaceiphila* | 0 | 0 | 0 | 0 | 1 | 1 | 0 | 1 | 0 |
| *P. atrosepticum* | 0 | 0 | 0 | 1 | 0 | 1 | 1 | 0 | 1 |
| *P. carotovorum* | 1 | 1 | 0 | 1 | 0 | 1 | 1 | 0 | 1 |
| *P. parmentieri* | 0 | 0 | 0 | 1 | 0 | 1 | 1 | 0 | 1 |
| *P. polaris* | 0 | 0 | 0 | 1 | 0 | 1 | 1 | 0 | 1 |

The strains used in this study are *D. dadantii* 3937, *D. aquatica* 174/2, *D. chrysanthemi* ATCC 11663, *D. dadantii* subsp *dieffenbachiae* NCPPB 2976, *D. dianthicola* NCPPB 453, *D. fangzhongdai* DSM 101947, *D. lacustris* S29, *D. oryzae* ZYY5, *D. paradisiaca* ATCC 33242, *D. poaceiphila* NCPPB 569, *D. solani* IPO 2222, *D. undicola* 2B12, *D. zeae* NCPPB 2538, *P. atrosepticum* ATCC 33260, *P. carotovorum* subsp. *carotovorum* ATCC 15713, *P. parmentieri* RNS08.42.1A and *P. polaris* NIBIO 1006. The presence and number of proteins detected by search of the corresponding gene in the genome in each strain is indicated.

**Table 2:**
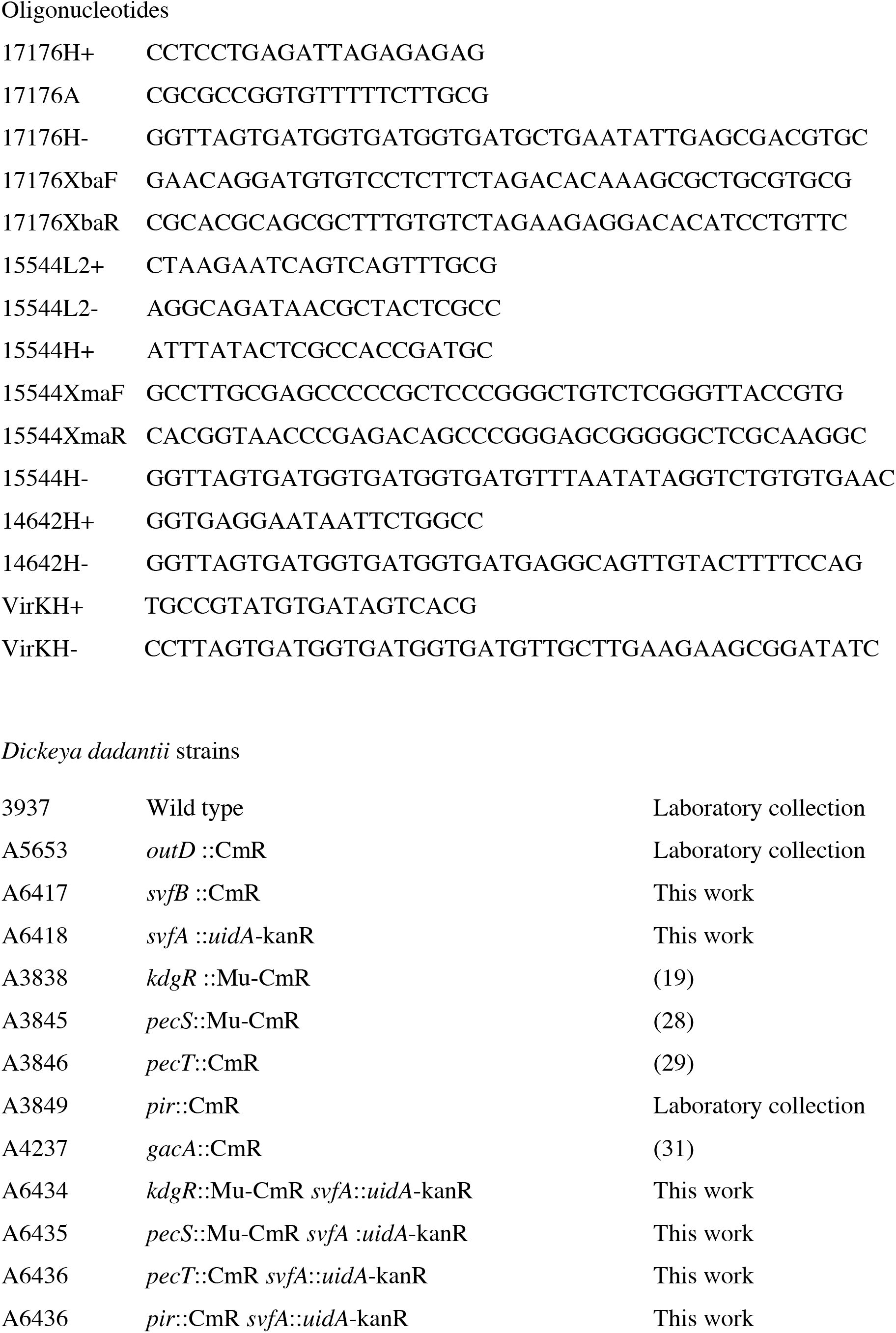
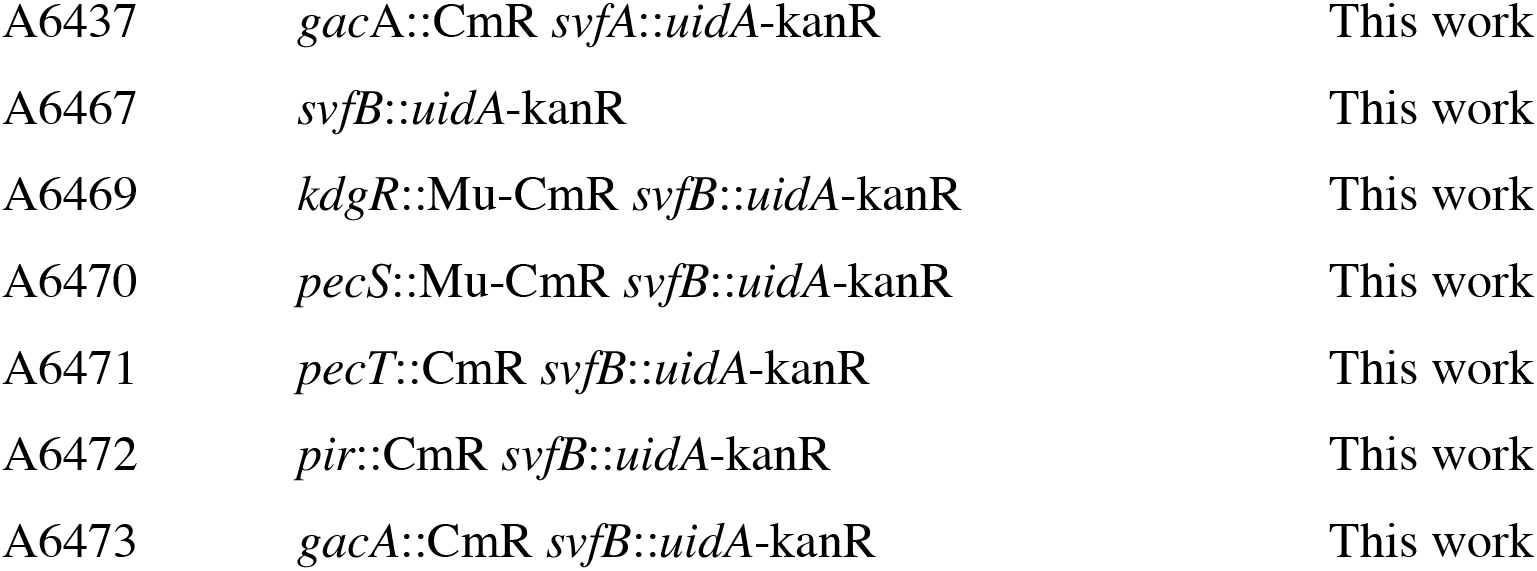
Oligonucleotides and strains used in this study.

We also examined the presence or absence of genes of other non-pectinolytic proteins known to be secreted by a T2SS in *Dickeya* or *Pectobacterium*: *ibpS*, *nipE*, *xynA*, *avrL*/*avrM* (Table 1). IbpS is a metal binding protein that prevents ROS-induced killing of bacteria (17). NipE is a toxin that provoke plant cell death (32). XynA is a xylanase that was identified in a corn strain of *Dickeya zeae* previously named *Erwinia chrysanthemi* (33). AvrL is homologous to the *Xanthomonas campestris* avirulence protein AvrL (15). Two very similar proteins, AvrL and AvrM, are encoded by *D. dadantii* 3937. AvrL was named Svx in *P. atrosepticum* where a role in virulence has been proven (34). However, its function in *Dickeya* has not been studied. IbpS is present in almost all the species, except *D. paradisiaca*. NipE is absent in *D. paradisiaca* and *D. poaceiphila*. Presence of XynA is variable in *Dickeya* strains and it is absent in *Pectobacterium*. A variation in the presence and number of AvrL can be observed in Dickeya strains (Table 1). Thus, the repertoire of T2SS-secreted protein known to be important for virulence is very variable from species to species.

## DISCUSSION

The T2SS of *Dickeya* and *Pectobacterium* is a major virulence factor of these bacteria. The knowledge of the repertoire of secreted proteins is necessary to better understand the precise mechanisms of virulence of these bacteria. These analyses have been undertaken with the model strain *D. dadantii* 3937 and partially with *Pectobacterium atrosepticum* (35)(16). *Dickeya* and *Pectobacterium* are characterized by their ability to degrade pectin and they are identified by this characteristic on the semi selective Crystal Violet Pectate medium. They all secrete enzymes capable of degrading pectin (pectate lyases, polygalacturonases, pectin methylesterases). However, recent works show that other proteins are secreted by the Out T2SS (15)(16). In the present work we used published transcriptome data to identify new potential substrates of the *D. dadantii* T2SS. The most highly induced genes in a transcriptome experiment of *D. dadantii* infecting *A. thaliana* are known virulence genes (*pelI*, *prtA*, *rhiE*, *paeY*, *rhaD*, *ibpS*, etc…) (19). However, in this top list some genes have no known function. The presence of a signal sequence in their product suggested that these proteins could be substrates of the T2SS necessary for the infection process. We showed here that the proteins SvfA, SvfB and YoaJ produced by genes present in the top list of those induced in Arabidopsis are substrates of the Out T2SS. YoaJ belongs to the family of expansins, proteins that loosen cellulose fibers. Their role as a virulence factor has been shown in *P. brasiliense* and *P. atrosepticum* (23) and it probably has the same function in *D. dadantii* and other *Dickeya* species. No function could be predicted for SvfA and SvfB which belong to the DUF 4879 family of proteins. However, a reduction of virulence of a *svfA* mutant and a *svfA svfB* double mutant on potato could be observed, proving a role of the proteins in the bacterial pathogenicity. Although an additive effect of the mutations was observed, they could not have exactly the same function. The mutants should be tested on various hosts to detect potential differences. It can be supposed that each protein would be more active on one type or one family of plant. Presence of three DUF 4879 proteins in *D. fanghzongdai* could explain its wide host range, from orchid to pear trees. Presence of homologues of SvfA and SvfB in *Photorhabdus* and in *B. thuringiensis* strains, two insect pathogens, indicates that the role of these proteins is not restricted to plant virulence but may participate to a common process of bacterial pathogeny. We also showed here that the PecS-regulated protein VirK is secreted by Out. No role on virulence had been observed for this protein with the chicory leaf model of infection (21). Other models should be tested to find the role of this protein.

Regulation of expression of the *svfA* and *svfB* genes is atypical for a *D. dadantii* gene involved in pathogeny. While expression of most of the virulence factors is induced in the presence of pectin or its derivatives through the repressor KdgR and repressed by glucose, that of *svfA* and *svfB* is opposite: it is activated by glucose and not controlled by KdgR. Expression of *svfA* is induced in the presence of plant tissue. This pattern of regulation has been described for *ibpS*, which is also strongly induced in *A. thaliana* (17). This could correspond to conditions encountered during the early phases of infection: pectin has not yet been degraded and glucose and saccharose are plentiful in plant tissues. *svfA* and *ibpS* could be among the earliest gene to be induced at the onset of infection, before the genes involved in pectin degradation. However, regulation of these genes by PecS and PecT shows that *svfA* and *svfB* are fully integrated in the network of regulators that controls *D. dadantii* virulence.

This work has extended our knowledge of the Out-dependent secretome of *D. dadantii*, showing that besides pectinases several other proteins are secreted. If the number of pectinolytic enzymes secreted is almost identical in the various *Dickeya* species, the number of additional secreted proteins varies markedly. Among the proteins analyzed (Table 1), *D. paradisiaca* has only one (VirK) while *D. fanghzongdai* has ten. All the intermediate combinations can be found in the various species. There seems to be less variations in the *Pectobacterium* strains surveyed. It is tempting to speculate that the presence/absence of these proteins could influence the host preference of some *Dickeya* species, providing additional virulence factors favorable to infect certain hosts. Works that compare *Dickeya* strains to understand what makes difference in their host range or aggressivity often focus only on the presence of the six known types of secretion systems without analyzing what proteins could be secreted (18)(36)(37). An exhaustive analysis of the secreted proteins would be more informative.

Is there other T2SS-secreted proteins to be identified in *Dickeya* strains? No specific signal is present on T2SS-secreted proteins that would allow their identification. 2D gels which were used in previous studies performed on *D. dadantii* and *P. atrosepticum* to identify their secretome have a limited sensitivity (15)(34). More sensitive methods such as liquid chromatography-tandem mass spectrometry (LC-MS/MS) can now be used (38). However, they give many false positive results since periplasmic and cytoplasmic proteins are often found in the culture supernatant. The approach we used here allowed the identification of four new secreted proteins. However, all these methods have a drawback. They can only detect proteins in conditions where they are produced. For instance, the rhamnogalacturonate lyase RhiE could only be detected when the bacteria were cultivated in the presence of rhamnose (39). The genes encoding YoaJ and VirK were not induced in *D. dianthicola* grown on potato (25). Another problem is that a protein may not exist in the strain tested. An analysis of the secretome of several *Dickeya* strains grown in several conditions will be necessary to have a global view of all the additional virulence factors that can be secreted by *Dickeya* species and evaluate their potential role in pathogenicity.

## ACKNOWLEDGMENTS

We thank Lison Massardier and Florence Ruaudel for technical work and Nicole Cotte-Pattat for reading the manuscript. This work was supported by funding from CNRS, University Lyon 1 and Agence Nationale de la Recherche (ANR-19-CE35-0016).

